# Non-muscle actinopathy-associated loss-of-function actin variants modulate cytoskeletal reorganization

**DOI:** 10.64898/2026.02.13.705838

**Authors:** Éva Gráczer, Kira Dakos, Tamás Bozó, Katalin Pászty, Nataliya Di Donato, Miklós Kellermayer, Andrea Varga

## Abstract

Variants in ACTB gene encoding for cytoplasmic β-actin result in a group of rare disorders called non-muscle actinopathies (NMA). We investigated the cellular effects of a missense variant, G302A, and a four-amino-acid deletion, S338-I341, associated with the subgroup of NMA – ACTB pLoF (predicted loss-of-function) disorder in patient-derived fibroblast cells. We found that neither of the mutations affected the organization of actin or the width of the actin-filament bundles, while the mutation G302A reduced the stiffness of the cells as measured by using atomic force microscopy. The latter effect might be associated with the misorganization of tubulin and with the increased size and number of focal adhesions. When we challenged the cells by monolayer stretching and followed the mechanically-induced reorganization of the actin cytoskeleton, we found that G302A mutant cells showed more dense actin filament bundles within the cells compared to wild type cells. At the same time, the extent of cofilin reorganization from the cell periphery was increased upon stretch, and this correlated with an increased cofilin phosphorylation. In the case of the deletion, while the extent of cofilin phosphorylation increased, the extent of reorganization was unaltered; rather, the phosphorylation of myosin light chain, important in counteracting external force, was drastically reduced. We could partially rescue this fascinating effect by overexpressing the active form of the formin mDia. Our findings open the possibility to validate the cellular phenotype in the most affected patient’s cells, in neurons.

## INTRODUCTION

Actin is a self-assembling biopolymer that plays a role in a wide array of physiological processes by forming dynamic networks: it contributes to maintaining the shape of cells, determines their mechanical properties and enables them to perform specific functions such as muscle contraction, cell motility and cell division [1]. The diverse cellular functions of actin are regulated by interactions with specific actin-binding proteins (ABP-s) [2]. Six actin isoforms are expressed in humans in a time- and tissue-specific manner. Four of them are muscle-specific (skeletal, cardiac, α- and γ-smooth) and two have cytoskeletal localization (β- and γ-cytoskeletal actin). The sequence identity of isoforms exceeds 93%, indicating that actin is a highly conserved intracellular protein [3, 4].

Mutations in genes ACTB and ACTG1 (encoding the β- and γ-cytoskeletal actin isoforms, respectively) cause a broad spectrum of rare disorders, called nonmuscle actinopathies (NMAs). NMAs show high clinical variability, which are classified into several groups (Figure 1A and [5]). Baraitser-Winter cerebrofrontofacial syndrome (BWCFF) patients are characterized by facial dysmorphism. This group of patients has the most severe symptoms, such as developmental delay, intellectual disability and cortical malformation associated with epilepsy [6]. The most frequent variant in BWCFF cohort is the substitution at the position R196 of cytoskeletal β-actin. The mutations of this residue have been connected to several defects. At the cellular level, patient-derived BWCFF-associated R196H fibroblasts showed reduced proliferation and migration rates, supporting the patient phenotype [7]. Actin stability was decreased through increased depolymerization and decreased polymerization rates [8]. The same mutation resulted in a decreased amount of F-actin in patient-derived R196H mutant fibroblasts [7]. In addition, this mutation also decreased the actin binding of several ABP-s, such as the Arp2/3 complex, cofilin and myosin 5A [7, 8]. Interestingly, cerebral organoids having either the mutation T120I in ACTB or theT203M in ACTG1 showed a reduction in size possibly due to the change in the mitotic cleavage plane angles [9]. The latter mutation was shown to decrease F-actin stability similarly to the R196H mutation [5]. There is a group of patients with mutations in either cytoskeletal actin genes categorized as “unspecified NMA” (Figure 1A). Among them, the effect of the cytoskeletal γ-actin carrying the mutation E334Q was shown to affect the interaction of actin with cofilin and myosin [10]. There is also a group of patients with mutations in the ACTG gene, that seems to be unaffected by the mutations. Another group of patients with mutations in the ACTG gene has hearing loss [11, 12]. A specific mutation in the ACTB gene, R183W, is exclusively connected to dystonia-deafness syndrome [13]. The last group of individuals (previously categorized into different subgroups) is relatively mildly affected and classified into a group named ACTB pLoF disorder. This group includes patients with syndromic thrombocytopenia and patients with ACTB-related pleiotropic developmental disorder [14–16]. The patients show mild intellectual disability and variable thrombocytopenia. Concerning the mutation landscape, point mutations, frameshift mutations or deletions affect the actin gene mostly in exons 5 and 6, although few frameshift or point mutations have also been found in exons 3 and 4 (Figure 1A). Frameshift (fs) mutations in exons 3 (L110Rfs, [15]) and 4 (V159Gfs, [5]) have been identified. Further mutations include point mutations in exon 5 (G302A, [14] and M313R, [16]), frameshift mutations in either exon 5 (T297Sfs, [15]) or 6 (A331Vfs, S368Lfs, [16]) and a deletion in exon 6 (S338-I341del, [16]). Two patients (mother and son) harbored an intrachromosomal deletion, which completely removed one allele of the ACTB gene [14].

**Figure 1.**
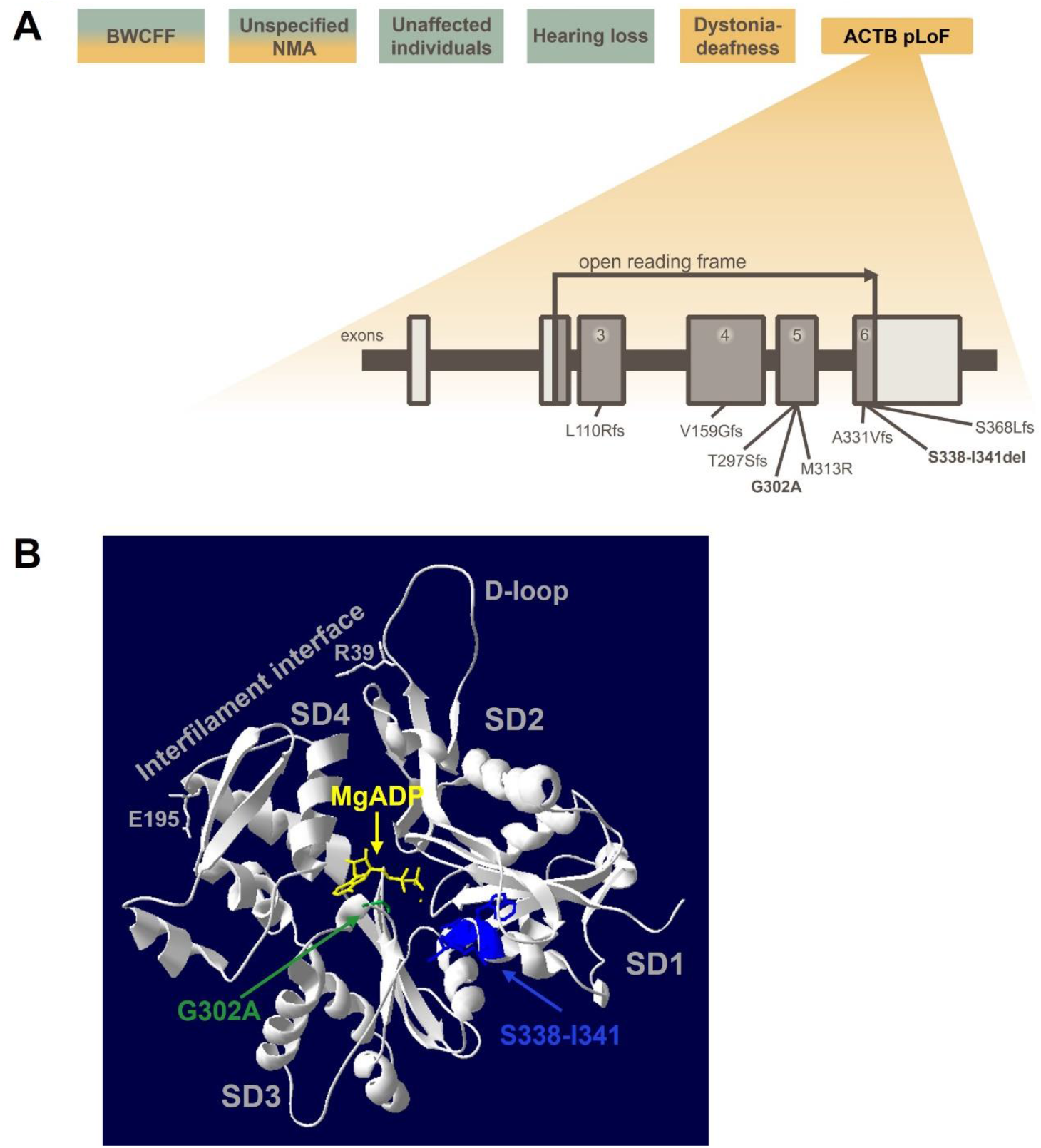
Location of the residues associated with ACTB pLoF disorder. **(A)** Classification of cytoskeletal actin mutations together with the schematic representations of mutations associated with ACTB pLoF disorder found in exons 3-6. Color code: symptoms associated with mutations in γ-actin or β-actin are labeled orange and green, respectively. BWCFF and unspecified NMA can be associated with both β- and γ-actin mutations. **(B)** Structure of wild type cytoskeletal β-actin (PDB code: 5JLH, [37]) is illustrated as grey ribbon structure, showing the location of the G302 residue (green) by stick model and the S338-I341 sequence (both the residues and the corresponding ribbon structure is colored as blue). The subdomains of actin are labeled as SD1-SD4 together with the D-loop and the interfilament interface (two residues are shown by stick models, which are involved in the interfilament interaction: R39 and E195). MgADP is shown by a yellow stick model.

In the study of Latham *et al*., among the ACTB pLoF variants the effect of A331Vfs or the S338-I341 deletion in patient-derived primary fibroblasts were compared to healthy control fibroblasts [16]. The mutant cells were smaller. Cells with S338-I341 deletion formed clusters, most probably due to stronger intercellular interactions [16]. In addition, mutant cells showed a decreased amount of β-actin in the lysates, while the amount of γ- and smooth-muscle actin were increased as a compensation for β-actin loss. Cell migration rate was reduced by both mutations, which is probably linked to the mild microcephaly phenotype. The recruitment of some ABP-s was also changed upon the mutations: increased expression of non-muscle myosin 2A (NM-2A), filamin A and α-actinin 1 was detected in the sub-nuclear region of the cells [16]. Molecular modeling of the interaction between actin and myosin in the presence of the S338-I341 deletion suggested significant changes in the interaction between the cardiomyopathy (CM)-loop of myosin and actin. Analysis of the platelets of the S338-I341 deletion variant showed increased non-muscle myosin 2 (NM-2) localization to the cortical actin bundles. The authors concluded that these mutations inhibit the final stages of platelet maturation through a mechanism affecting microtubule organization. It has been shown that the S338-I341 deletion reduced the expression of β-actin in insect cells, providing a hint for a less stable protein [5]. G302A [5] and S368Lfs mutations [17] decreased the thermal stability of recombinant β-actin, pointing at an allosteric perturbation of the actin structure. The effect of the G302A variant was investigated in blood mononuclear cells [14]. Here, the authors hypothesized that this variant disturbs the organization of microtubules, as they found a highly acetylated, lower molecular weight tubulin in the patient with the G302A mutation. From fractionation experiments the mutant blood mononuclear cells contained less β-actin in the detergent insoluble fraction, indicating that the mutant actin is less associated with the cytoskeleton.

Residue G302 is located at the nucleotide binding site of actin (Figure 1B, green), while the residues of the S338-I341 deletion are positioned at the beginning of a helix in subdomain 1 (SD1, Figure 1B, blue). The binding sites of the actin-binding proteins (ABPs) cofilin and myosin are located on the outer surface of the actin filament, involving residues from two subdomains, SD1 and SD2. The positions of both mutations are far from the interfilament interface. Based on the known structural data one may predict that mutation of the G302 residue possibly influences the binding of the nucleotide, which might also be reflected in the binding of cofilin [18, 19], known to be sensitive to the nucleotide-binding state of actin (preference for ADP-bound actin). The deletion, based on its location, might affect both cofilin and myosin binding. Based on these assumptions, actin reorganization might also be perturbed in the cells. The question arises whether the clinical phenotype can be connected to a specific actin-ABP interaction or lies in the perturbed dynamic organization of actin.

Here we report a thorough analysis of how heterozygous point mutation G302A and the deletion of the sequence S338-I341 in actin affect the behavior and properties of patient-derived dermal fibroblasts. In addition to the analysis of the actin organization, we compared cell stiffness with atomic force spectroscopy. Reorganization of actin was investigated by monolayer stretching. During actin reorganization we followed the localization of the actin depolymerizing factor cofilin. We also monitored the phosphorylation of cofilin (required for its inactivation and its dissociation from actin) and myosin light chain (MLC), as indicators of the efficiency of actin contraction upon external stretching forces. We find that both mutations affect actin reorganization upon applied force, but in different ways: while the G302A mutation affects the cofilin-mediated step, the deletion disturbs the contraction necessary to withstand the external force.

## RESULTS

### G302A mutation impairs cell stiffness and tubulin organization in patient-derived fibroblasts

Patients with the A331Vfs mutation or the S338-I341 deletion show mild microcephaly, which is supported by the migrational defect of patient-derived fibroblasts [16]. By contrast, a patient with the mutation G302A showed a borderline microcephaly only [14]. To investigate whether this milder phenotype can be recapitulated in patient-derived fibroblast cells, we compared cell growth (Figure 2A) and cell migration (Figure 2B and Figure S1A,B) of wild type and G302A fibroblasts. We found no significant differences either in cell growth or migration. Since the patient has mild intellectual disability and attention deficit, we wanted to find out whether this can be connected to any disturbed cellular property of actin. Therefore, we investigated the cellular organization of actin. First, we compared the localization of actin by phalloidin staining in wild type, G302A and S338-I341del patient-derived fibroblast cells (Figure 2C). We did not observe any striking differences in phalloidin staining, but the smaller cells obtained upon the mutation S338-I341del showed less pronounced zonula occludens (ZO-1) staining, indicating that tight junctions were less pronounced. Then, we investigated the structure of the actin filament bundles by superresolution (STED) microscopy (Figure 2D) and quantified the filament width from the fit (Figure 2E) of the Gaussian cross-sectional intensity profile (Figure S1C,D). Although the width of the filament bundles was slightly reduced in both mutants, the differences were not significant (Figure 2E). Based on the width of the actin double helix (7-9 nm) [20], the estimated number of filaments was 15-20 in the wild type and 14-18 in the mutant cells. To uncover whether the mutations have any effect on cell stiffness, we carried out atomic force microscopy on live fibroblasts. Typical force curves for wild type (WT), G302A and S338-I341del cells are shown in Figure 3A. Distribution of the elastic moduli of wild type and S338-I341del mutant cells (Figure 3B) revealed that there is a small difference in the full width at half-maximum (FWHM), but the mean value of the distribution (Figure 3C) was not affected by the mutation. However, for G302A the mean value of the elastic modulus dropped to about one fourth of that in the wild type cells. This indicates that the G302A mutation reduces cell stiffness considerably, which cannot be accounted for by the small change in actin bundle width. To uncover the mechanisms behind the reduced stiffness, we compared the amount of phosphorylated myosin light chain (pMLC) by western blotting in wild type and G302A cells (Figure 3D). We found no significant differences in the extent of MLC phosphorylation. Thus, the reduced elastic modulus of the G302A mutant cells (Figure 3C) cannot be explained by a decrease in contractility. As in the case of the S338-I341del mutant a reduced amount of β-actin was detected in cell lysates [16], we also checked whether the G302A mutation affects the amount of β-actin. We quantified the amount of β-actin and pan-actin of the cell lysates of wild type and G302A fibroblasts (Figure 3E), but we found no significant differences between the two cohorts. To investigate whether a reduced F-actin content of the G302A cells can explain the reduced cell stiffness, we quantified the amount of F-actin in wild type and G302A cells. The amounts of F-actin and G-actin separated by ultracentrifugation showed no significant difference (Figure 3F) between the wild type and the G302A mutant cells as analyzed by either a β-actin or a pan-actin antibody. Since the organization of tubulin might also account for the changed stiffness, we compared the localization of tubulin both in wild type and G302A cells (Figure 3G). We found an increased perinuclear localization of tubulin in the G302A mutant cells. To confirm that these changes are indeed related to tubulin depolymerization, we treated wild type cells with different concentrations of colchicine (Figure 4A). Upon increasing colchicine concentration, the fluorescence intensity of tubulin in the perinuclear region was increased, while the microtubular network formation was reduced, which was the most obvious at the highest concentration of colchicine. This phenomenon was quite similar to that in G302A mutant cells without colchicine treatment. Since microtubule depolymerization promotes focal adhesion growth [21], we analyzed the focal adhesions in wild type and G302A mutant fibroblasts (Figure 4B) by staining for focal adhesion kinase (FAK) phosphorylated on Tyr397 (pFAK). We found that the focal adhesion number (Figure 4C) and size (Figure 4D) was significantly increased by the G302A mutation.

**Figure 2.**
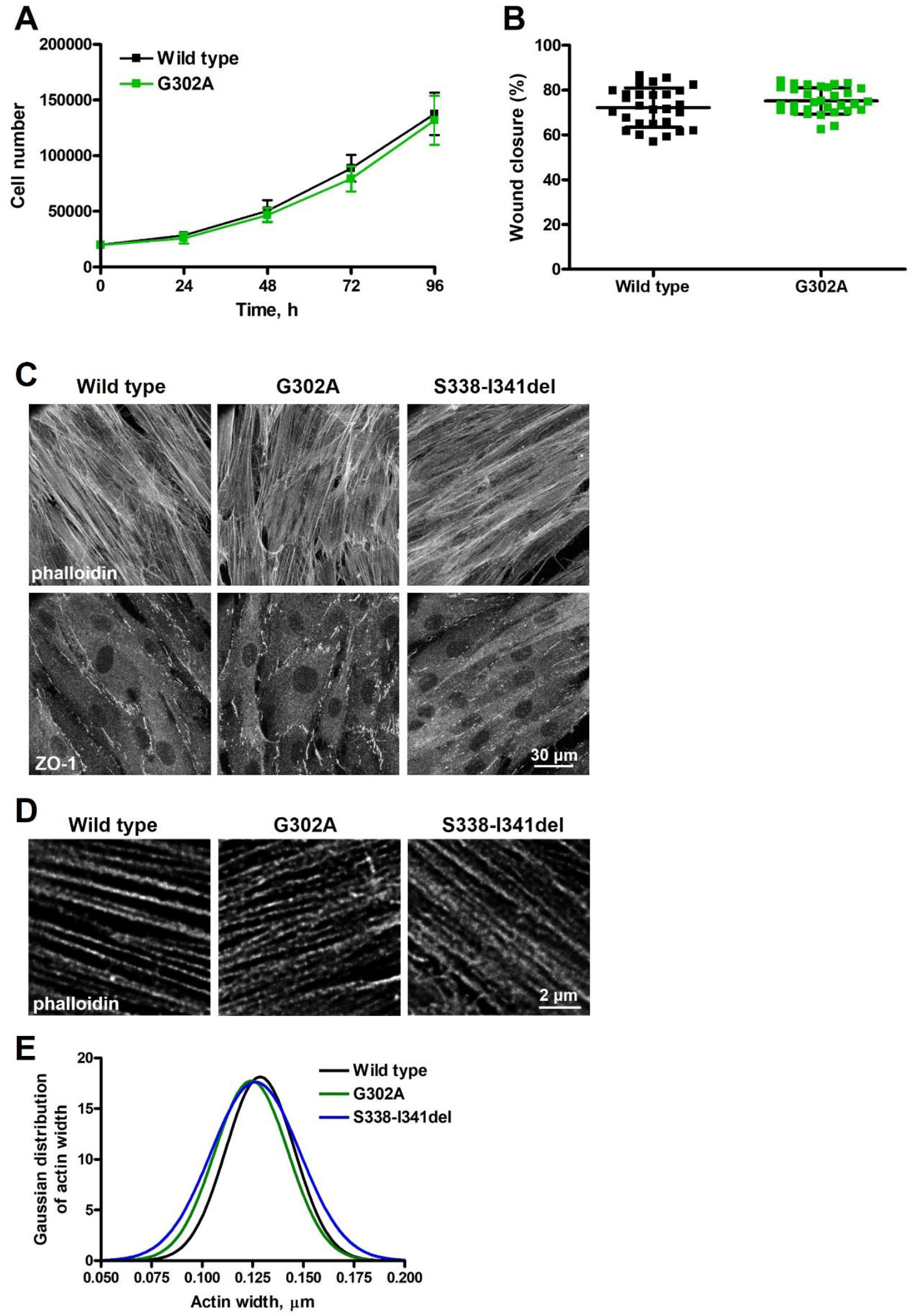
ACTB pLoF disorder - associated mutations do not impair actin filament organization in patient-derived fibroblasts. **(A)** Cell growth curves of wild type (black squares) and G302A (green squares) cells are shown. Data are plotted as mean ± SD. **(B)** Cell migration was monitored in a wound closure assay (Figure S1A,B). The percentage of wound closure was plotted for wild type (black squares) and G302A (green squares) cells. Data shown in (A) and (B) are from three independent experiments. **(C)** Confocal images of fixed wild type, G302A mutant and S338-I341del fibroblasts stained with phalloidin and zonula occludens (ZO-1). **(D)** Superresolution STED images of wild type, G302A mutant and S338-I341del cells stained with phalloidin. **(E)** Gaussian fit of the distribution profile of actin width quantified from the line profile after image deconvolution (Pairwise comparison of the histograms of wild type as well as of the mutants are shown in Figure S1C,D). Data shown in (C) and (D) are from two independent experiments.

**Figure 3.**
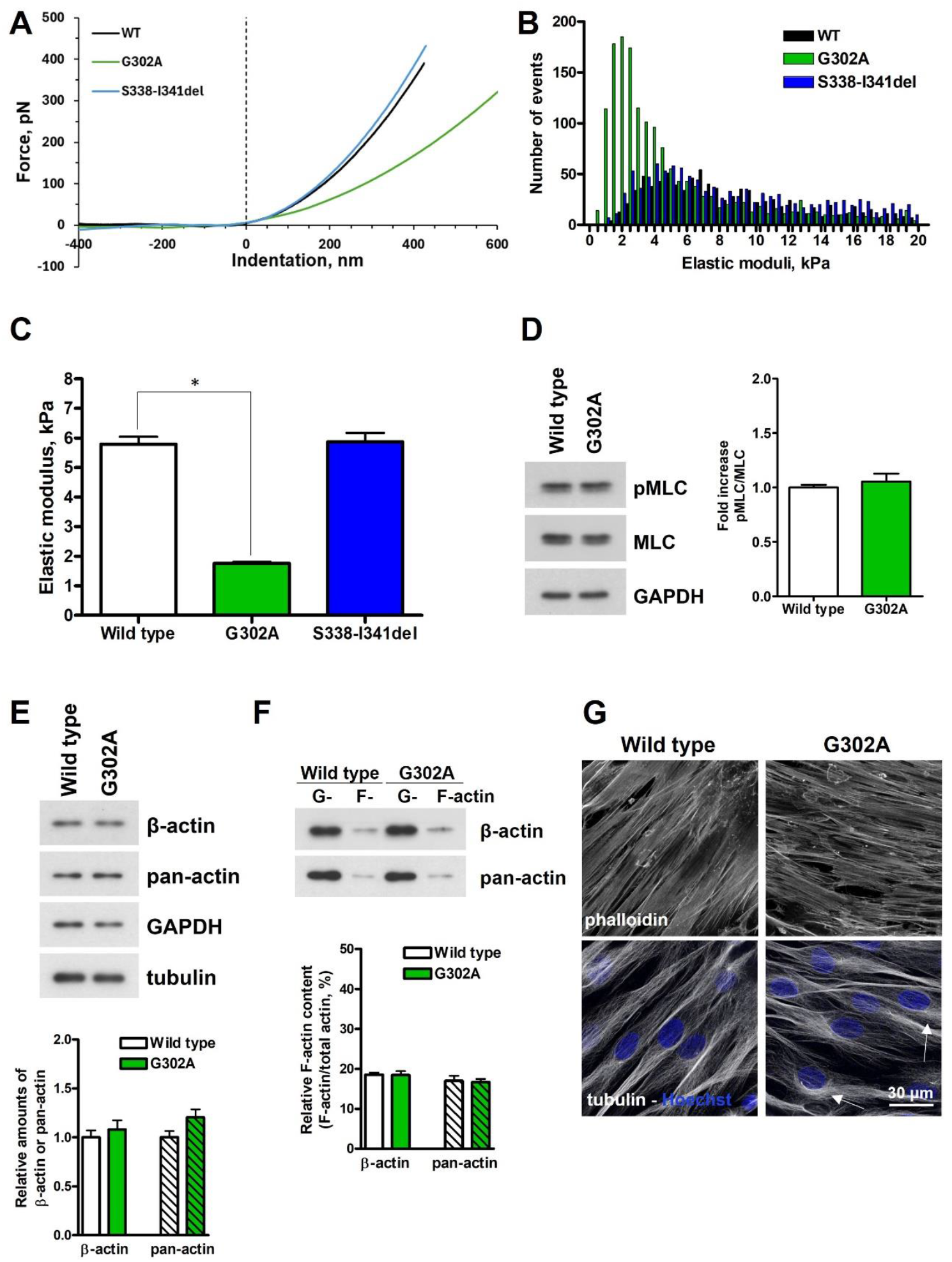
G302A mutation decreases the cellular stiffness of patient-derived fibroblasts. **(A)** Representative force-indentation curves of live wild type (WT, black), G302A mutant (green) and S338-I341del fibroblasts (blue). **(B)** Distribution of the elastic moduli of wild type (WT, black), G302A (green) and S338-I341del cells (blue) calculated from the force-distance curves as described in the Materials and Methods section. **(C)** Mean values of the Gaussian fit of the histograms (calculated from the log-normal distribution) are shown for wild type (black), G302A (green) and S338-I341del cells (blue). **(D)** Phosphorylation of MLC was compared in wild type and G302A cells by western blotting. Quantification of the amount of pMLC (normalized to MLC) is plotted on the right. **(E)** Total lysates of wild type and G302A mutant fibroblasts were immunoblotted and labeled with the antibodies of β- and pan-actin. Tubulin and GAPDH were also labeled as loading controls. Results of two independent experiments are shown. Quantification shows the relative amount of β-actin or pan-actin normalized to the amount of GAPDH and plotted below. **(F)** F-actin (recovered from the pellet after ultracentrifugation) and G-actin (supernatant) content of wild type and G302A mutant fibroblast cells were determined by western blotting using a β-actin antibody or a pan-actin antibody. The amount of F-actin was quantified for each condition as the ratio of F-actin and the total actin (the sum of F- and G-actin) for each sample separately and it is plotted below. **(G)** Confocal images of fixed wild type, G302A mutant cells stained with phalloidin, tubulin and Hoechst. The lower panel set shows the merged image of the tubulin (grey) and Hoechst (blue) stainings of wild type and G302A cells. White arrows in panel E indicate examples of the increased perinuclear staining in the mutant (G302A) cells. Data are from two independent experiments. Data were analyzed by one-way ANOVA, followed by Bonferroni’s multiple comparison test expressed as mean ± SD (*p < 0.05).

**Figure 4.**
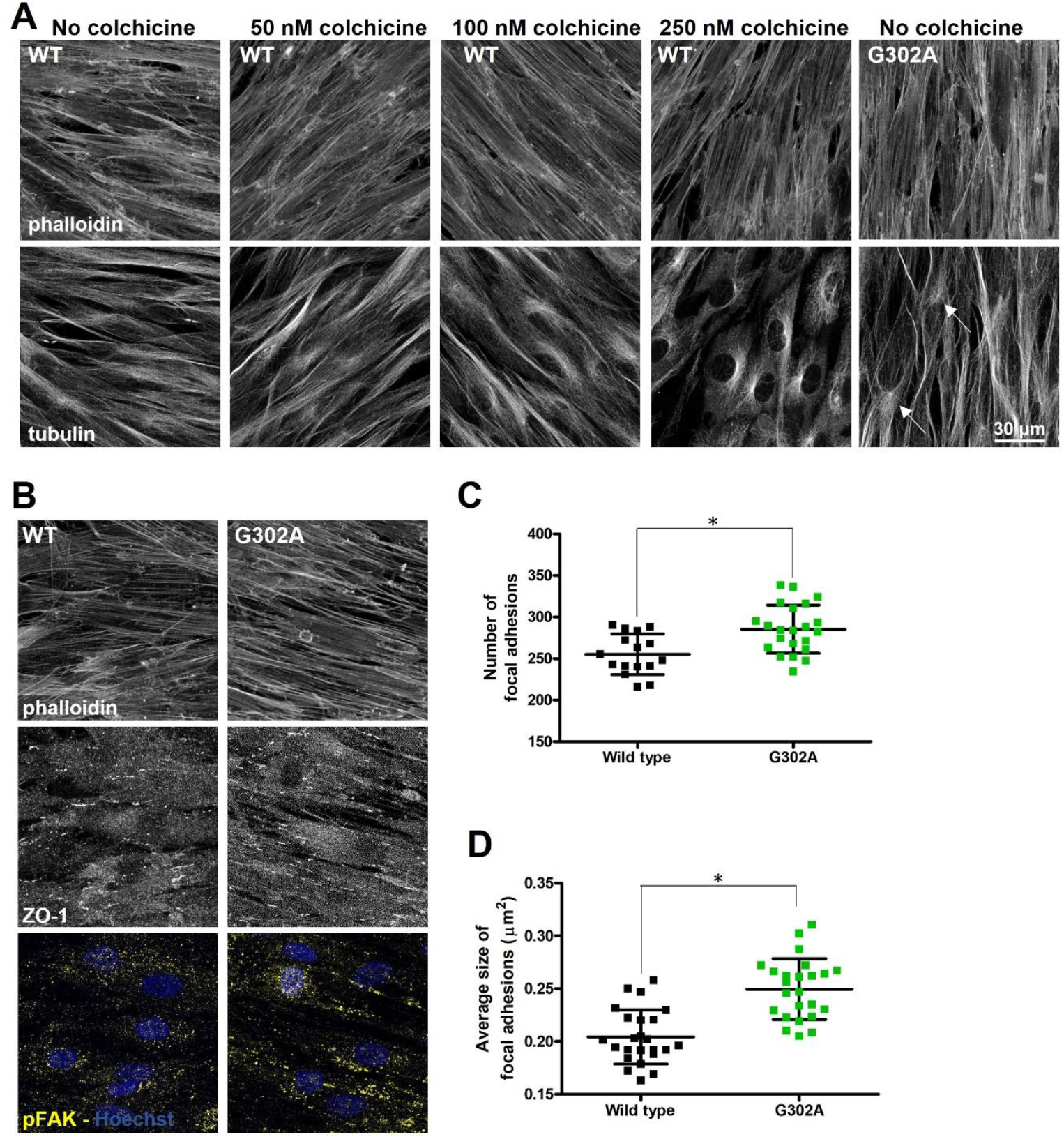
G302A mutation perturbs the organization of the microtubules and focal adhesions of patient-derived fibroblasts. **(A)** Confocal images of wild type (WT) or G302A cells treated with different concentrations of colchicine (or ethanol) for 1 hour. “No colchicine stands” for cells treated with ethanol, what was used to dissolve colchicine. Fixed cells were stained with phalloidin and tubulin. **(B)** Wild type (WT) and G302A cells were fixed and stained with phalloidin, ZO-1 and pFAK antibodies, and Hoechst. The lowest panel shows the merged images of pFAK (yellow) and Hoechst (blue) stainings. **(C)** Quantification of the number of focal adhesions (pFAK-labeled spots) determined in a 140 x 140 μm area of wild type (black filled squares) and G302A (green filled squares) cells are shown. **(D)** Quantification of the average size (area) of focal adhesions (pFAK-labeled spots) determined in a 140 x 140 μm area of wild type (black filled squares) and G302A (green filled squares) cells are shown.

### G302A mutation impairs cofilin reorganization, while S338-I341del mutation decreases myosin light chain phosphorylation upon stretch

To find out whether the investigated mutations affect the reorganization capacity of actin, we stretched wild type and mutant cells by 30% of their length, kept them stretched for 15 minutes, then fixed and stained for phalloidin and cofilin (Figure 5A). By comparing phalloidin staining, we could observe more actin fibers within the cells in the stretch direction in G302A mutant fibroblasts, whereas in the case of the S338-I341del mutation actin staining was stronger near the cell-cell junctions, but not in the cell’s interior. Cofilin removal from the cell periphery upon stretch was increased only in the case of the G302A mutant relative to its wild type counterpart (Figure 5B). To correlate the extent of changes in cofilin reorganization with cofilin phosphorylation (which enhances its dissociation from actin), we terminated the stretch at different time points by cell lysis to reveal the kinetics of cofilin phosphorylation by western blotting (Figure 6A and B). Cofilin phosphorylation was slightly increased already in the absence of stretch and remained constant in the G302A mutant (Figure 6B). This correlates with the increased extent of cofilin dissociation from actin (Figure 5B). While in the case of the S338-I341del mutant we also observed an increased cofilin phosphorylation (Figure 6C and D), it was not accompanied by an increased cofilin dissociation (Figure 5B). As a result of the uniaxial stretch, parallel actin bundles are formed in the direction of stretch and those might be decorated by myosin so that the cells can counteract the external force in the direction of the stretch. By monitoring the extent of myosin light chain phosphorylation (Figure 6E and F) we found no major difference between wild type and G302A mutant cells, while there was a drastic reduction of myosin light chain phosphorylation upon the S338-I341 deletion (Figure 6G and H). This was already observed before stretch and although the extent of phosphorylation increased, it never reached the level of that in the wild type cells. It was shown that myosin II cooperates with formin (responsible for linear actin cable formation) in the regulation of force generation [22]. Therefore, we attempted to rescue the myosin light chain phosphorylation defect observed in the S338-I341del cells by overexpressing the active form of the formin, mDia (Figure S2A-C). Western blot analysis of the lysates (Figure 7A and B) indicated a partial rescue of myosin light chain phosphorylation both in non-stretched (0’) and stretched (20’) cells.

**Figure 5.**
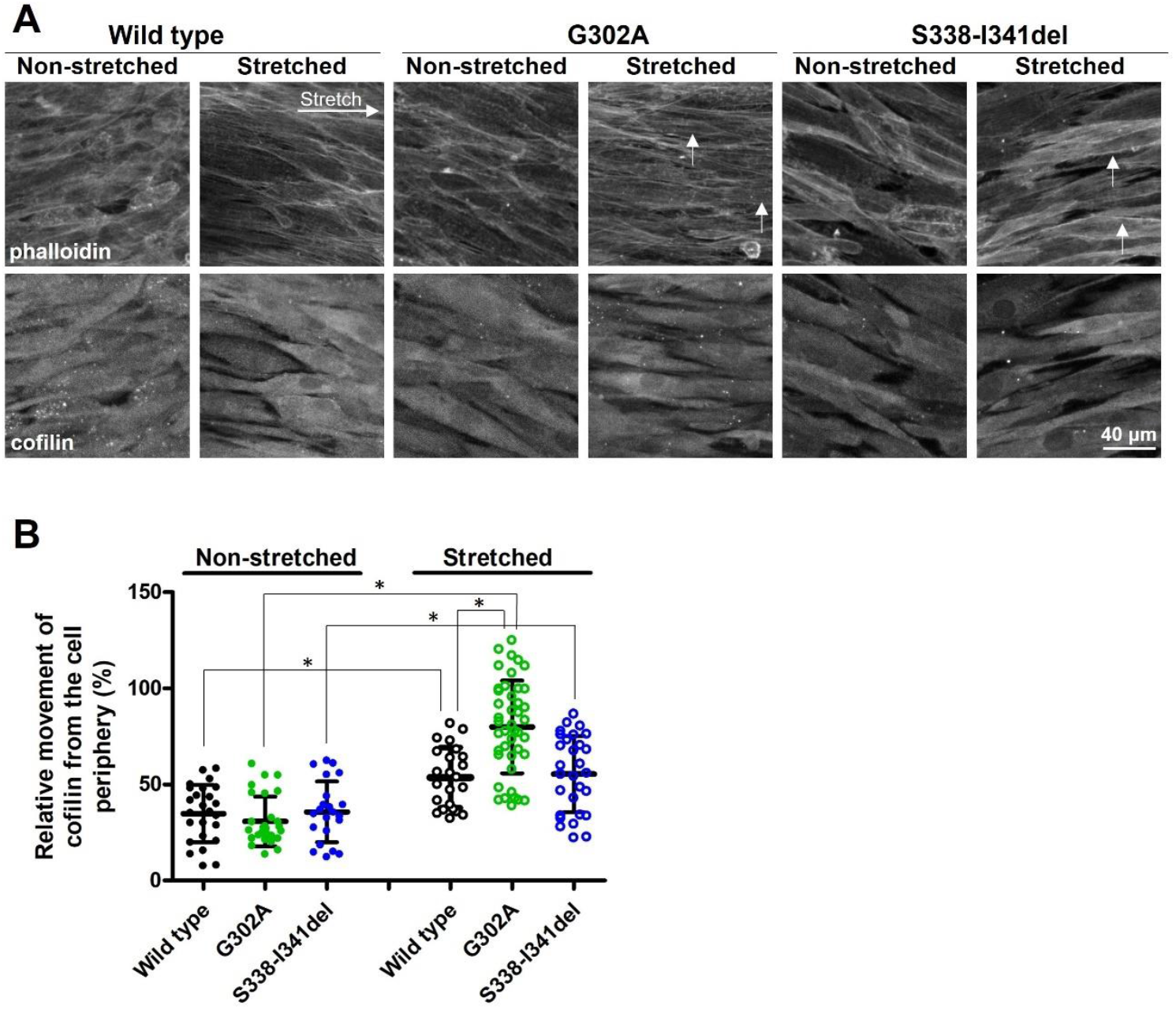
G302A mutation increases the dynamic reorganization of actin by enhancing cofilin dissociation. **(A)** Immunofluorescence analysis of wild type, G302A mutant and S338-I341del fibroblasts kept either non-stretched or stretched for 15 minutes. Samples were stained with either phalloidin or for cofilin. White arrows indicate the ticker internal stress fibers for the G302A mutants and the ticker peripheral actin bundles for the S338-I341del mutant. **(B)** Quantification of the relative movement of cofilin from the cell periphery. Non-stretched samples of wild type cells are shown in black, filled circles, while stretched samples are represented by black open circles. Samples of G302A cells are labeled as green, samples of S338-I341del fibroblasts are labeled as blue filled and open circles for non-stretched and stretched samples, respectively. Data are from two independent experiments. Data were analyzed by two-way ANOVA, followed by Bonferroni’s multiple comparison test expressed as mean ± SD (*p < 0.05).

**Figure 6.**
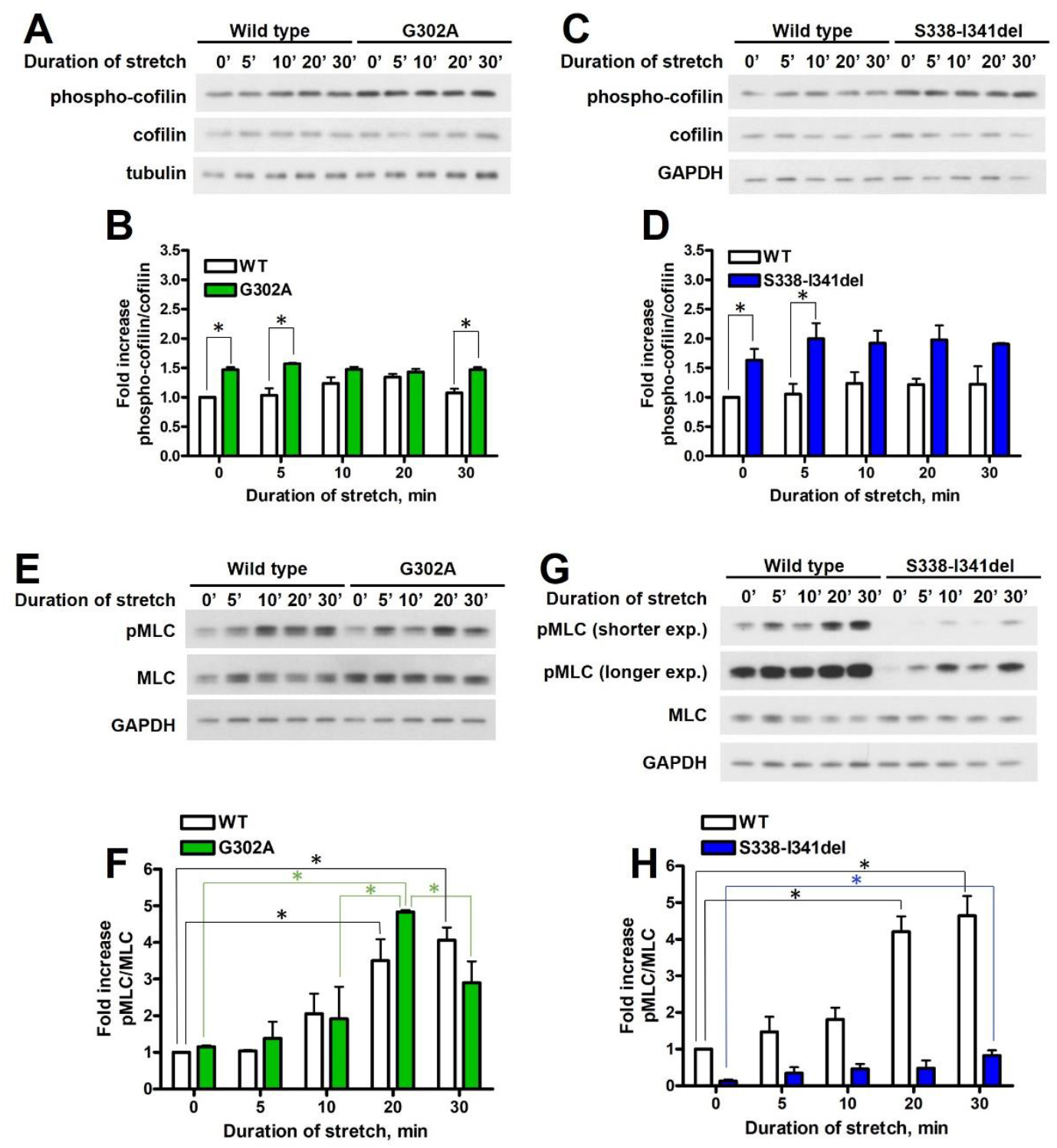
S338-I341 deletion impairs myosin light chain phosphorylation. **(A)** and **(E)** Wild type or G302A fibroblasts were stretched by 30% of their original length and were harvested at different time points. Phosphorylation of cofilin and myosin light chain (MLC) was analyzed by Western blotting. Quantification of the data is shown in **(B)** for p-cofilin and in **(F)** for pMLC. **(C)** and **(G)** Wild type or S338-I341del fibroblasts were stretched by 30% of their original length and were harvested at different time points. Quantification of the data is shown in **(D)** for p-cofilin and in **(H)** for pMLC. The amounts of pMLC and p-cofilin were normalized to the total MLC or cofilin amounts, respectively. Data are from two independent experiments. Mean ± SD is plotted. Data were analyzed by two-way ANOVA, followed by Bonferroni’s multiple comparison test expressed as mean ± SD (*p < 0.05).

**Figure 7.**
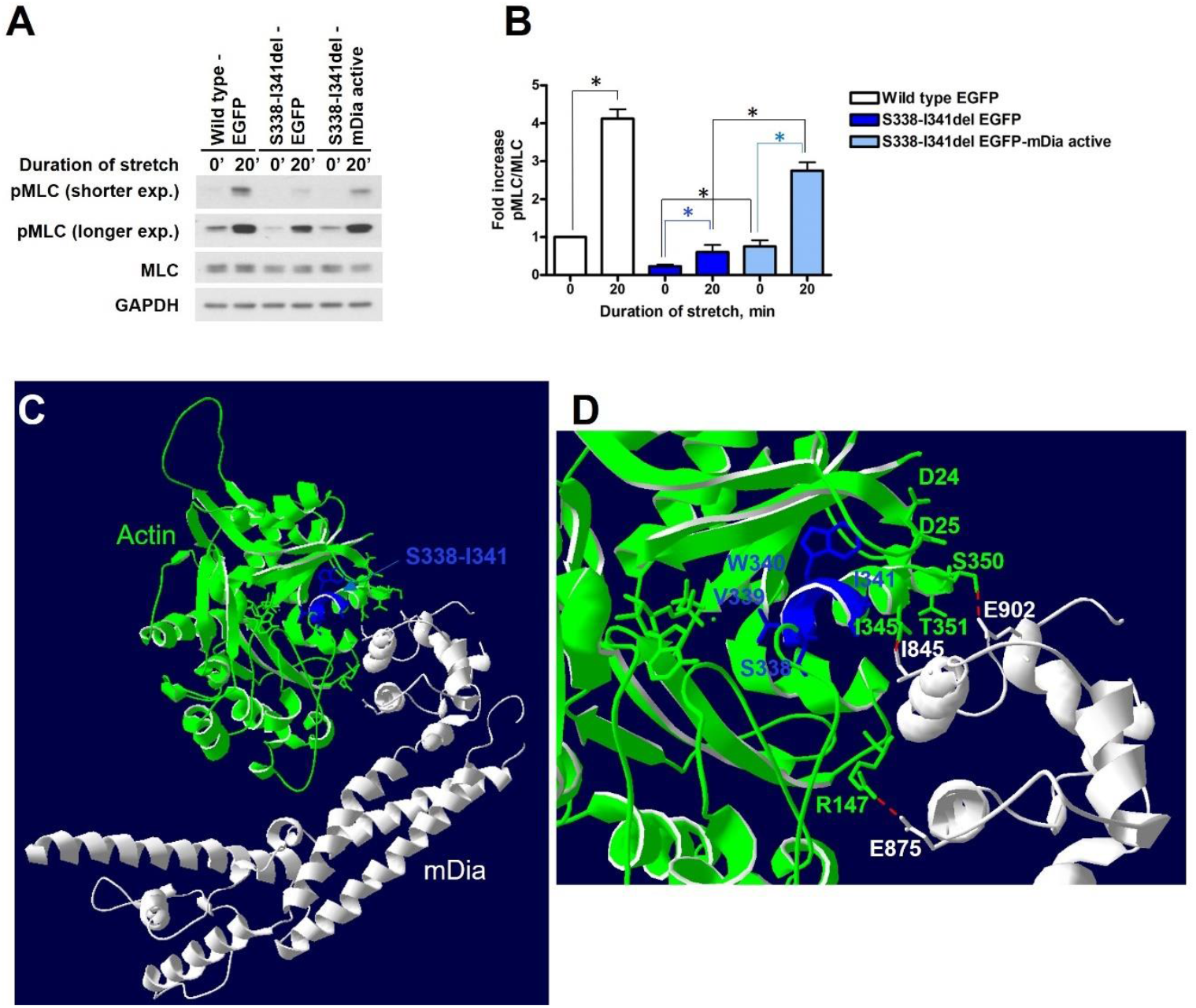
Defect in myosin light chain phosphorylation of S338-I341del actin can be rescued partially by overexpressing mDia. **(A)** Western blot analysis of wild type fibroblasts infected with EGFP, S338-I341del fibroblasts infected with EGFP, and S338-I341del fibroblast infected with EGFP-mDia active left either unstretched or stretched for 20 minutes before cell lysis. **(B)** Phosphorylation of myosin light chain was quantified and normalized to the total MLC amount. Data are from two independent experiments. Mean ± SD is plotted. Data were analyzed by two-way ANOVA, followed by Bonferroni’s multiple comparison test expressed as mean ± SD (*p < 0.05). **(C)** mDia (part of the FH2 domain, residues 832-1147) binding to actin as represented by the recently published cryo-EM structure (PDB code: 8RU2, [48]). mDia (grey) binding to an actin subunit (green) is represented on the figure, showing the S338-I341 segment (in blue). Residues interacting with the FH2 domain of mDia are represented by stick models. **(D)** Zoomed-in view of the interacting surface highlighting the possible interactions between the residues of actin and mDia are shown as red dashed lines.

## DISCUSSION

ACTB pLoF disorder has less severe clinical consequences that BWCFF [14, 16]. Very little is known about how the associated actin mutations impair the function of actin and of the cell altogether. We did not find major differences either in actin organization or the size of the actin bundles in patient-derived fibroblast cells. However, in the case of the G302A mutant we observed a decreased cellular stiffness, which is unlikely to be associated with changes in either the organization or the contractility of the actin cytoskeleton. Although in mononuclear blood cells there was an indication that the mutant actin has a decreased ability to form F-actin [14], we did not observe such in patient-derived fibroblasts, as revealed by the relative quantities of G- and F-actin. Thus, the reduced cell stiffness might be linked to how actin regulates the localization and function of microtubules. Indeed, the application of the tubulin depolymerizing drug, colchicine [23] in wild type cells showed a similar reorganization of the tubulin cytoskeleton as it was observed in the G302A mutant ones without the drug. Thus, the G302A mutation might influence the stability of microtubules. It has been shown that focal adhesion disassembly is regulated by microtubule growth [24] and, in turn, microtubule depolymerization promotes focal adhesion growth [21]. Indeed, we have found that in the G302A fibroblasts both the size and the number of focal adhesions were increased, as identified by the autophosphorylated form (pTyr397) of focal adhesion kinase (FAK) [25]. To explain the findings we hypothesize that the protein KANK, identified as a linker between the actin-binding protein (ABP) talin and the microtubules [26, 27], may be involved. KANK knockdown was reported to lead to the depolymerization of microtubules and the subsequent release of the Rho GEF, GEF-H1 from microtubules. Upon GEF-H1 release, Rho activity increased, and through the subsequent activation of ROCK, the increased phosphorylation of myosin light chain [28] might explain the increased growth of focal adhesions. Thus, it might be possible that the G302A mutation weakens the interaction of actin with microtubules through the interacting proteins (such as KANK) and this uncouples the microtubules from actin, leading to the depolymerization of microtubules.

Whether tubulin misorganization could play a role in the patient’s phenotype still needs to be addressed in neurons. It is clear, however, that the interplay between the actin cytoskeleton and the microtubules is important for neurite outgrowth. The dynamic reorganization of actin is necessary for creating protrusions with stable microtubules to determine the polarity of the axon [29]. Dendrites are also developed from such protrusions, but dendrites harbor microtubules with mixed polarity. It was shown that inhibition of microtubule polymerization might affect dendritic branching [30], and the extent of branching correlates with the level of intelligence, i.e. lower branching might be associated with intellectual disability [31]. This might be connected with the moderate phenotype of the ACTB pLoF patients [32].

The common feature of the investigated mutations is that both of them affect the reorganization properties of actin during uniaxial stretch. Actin reorganization is regulated by the actin depolymerizing factor cofilin [33]. As we observed previously, in endothelial cells cofilin becomes phosphorylated upon stretch [34]. During stretch-induced actin reorganization cofilin is dissociated from peripheral actin. Therefore, a change in cofilin localization can be observed at the cell periphery upon stretch. This localization difference between non-stretched and stretched cells can be quantified in both wild type and mutant cells. We found that the extent of cofilin “removal” from the cell periphery was increased in the case of the G302A mutant compared to the wild type fibroblasts. This might indicate that the binding of cofilin to the G302A mutant actin is decreased during actin reorganization. In line with this observation, we found a slight increase in cofilin phosphorylation in both the absence and presence of stretch.

In the case of the deletion variant, we did not find any differences in cofilin localization. The extent of myosin light chain (MLC) phosphorylation, however, decreased in the mutant, suggesting that this mutation affects the contraction efficiency of actin fibers upon mechanical stress. MLC phosphorylation has been shown to increase actin binding to myosin [35]. Therefore, the question arises whether myosin binding might be weakened, or whether the interaction of actin with myosin is changed upon binding to the mutant actin in such a way that affects the availability of the light chain to be phosphorylated by Rho-associated kinase (ROCK) or myosin light chain kinase (MLCK) [36]. By analyzing the available structural data, we found that the myosin binding site is close to the helix, the beginning of which contains the S338-I341 segment (labeled blue in Figure S3A,B). This segment does not interact directly with myosin but might help the positioning of a loop consisting of the residues D24 and D25 (through the interaction of W340 and P27). This loop interacts with the cardiomyopathy (CM) loop of myosin (Figure S3A,B [37]). In addition, the loop following the helix with the deletion contains two residues, S350 and T351, which interact with E556 of myosin. The mutation of the latter residue in *Dictyostelium discoideum* myosin (residue E531) has been shown to reduce the binding of myosin to F-actin to one tenth [38]. Thus, the deletion might perturb the positioning of the residues S350 and T351, which in turn could reduce myosin binding through the residue E556 to S338-I341del actin.

There are no available treatment options for the patients with ACTB pLoF disorder so far. The question arises whether it might be possible to devise a therapy based on the specific actin-ABP interactions changed by the mutations. In the case of the deletion variant the interaction of actin with myosin II was perturbed already in resting cells, and this reduction was also reflected in the MLC phosphorylation in response to stretch. A recent study raised the possibility of a crosstalk between myosin II and formin functions in the regulation of force generation [22]. The authors found that traction forces exerted by the stress fibers depend not only on myosin II, but also on formin. Therefore, we attempted to rescue the cell-contraction defect by overexpressing the active form of the formin, mDia. The overexpression partially restored MLC phosphorylation in the mutant, which indeed points at the crosstalk between formins and myosins, which results in adjusting the extent and duration of contraction. We also checked the binding site of the FH2-domain of formins on actin based on the available structural data. Indeed, the FH2 domain interacts with the surrounding of the helix with the deletion (I345 from the helix having the deletion, and S350, right after the helix, Figure 7C and D) and might potentially help to restore myosin binding.

This contractility defect observed in patient-derived fibroblasts might be relevant in the phenotype of the patients, since it has been shown that mutations in myosin II are linked to neurodevelopmental and neurodegenerative disorders [39]. Mutations associated with non-muscle myosin IIA are specifically linked to autism, intellectual disability and schizophrenia [36]. Migration of neurons during brain development is a critical event to establish appropriate connections between them [40]. Pharmacological inhibition of myosin activity showed the essential function of myosin in neuronal migration as it disrupted the forward movement of cells, and the pulling forces necessary for translocating the soma during neuronal migration [41]. Thus, the migration defect found in the S338-I341del cells [16] might be explained by the defect in myosin phosphorylation and contraction.

Taken together, we found that two variants associated with ACTB pLoF disorder affect the reorganization properties of actin, which might be important for neuronal migration, necessary for creating a properly connected neural network [42]. An interesting additional observation was in the case of the G302A mutant that the structural organization of tubulin was affected by the actin mutation. This, indeed, might have also critical consequences in neurons [30], affecting the branching ability of the cells. As no treatment is available for these patients so far, it would be critical to determine to what extent these defects appear in neurons and whether the proposed perturbation to rescue the defect in cell contraction would be a possible treatment option for the patients having the S338-I341 deletion in β-actin.

## MATERIALS AND METHODS

### Primary antibodies for western blotting (WB)

β-actin antibody, Cat# MCA5775GA from Bio-Rad; pan-actin antibody, clone 2A3 Cat# MABT1333 from Sigma Aldrich; GAPDH antibody Cat# 97166; MLC antibody, Cat# 8505S; pMLC (Thr18/Ser19) antibody, Cat# 3674S; cofilin antibody, Cat# 5175S; p-cofilin antibody, Cat# 3313S; all from Cell Signaling Technology; tubulin antibody, Cat# PA5-58711, from Thermo Scientific; anti-GFP antibody, Cat# GFP-1020 from Aves Labs.

### Primary antibodies for immunofluorescence (IF)

cofilin, Cat# ab42824 from Abcam, tubulin antibody, Cat# PA5-58711, from Thermo Scientific. ZO-1, Cat# 33-9100 and pFAK (Tyr397), Cat# 44-624G, both from Invitrogen.

### Secondary antibodies for WB

Peroxidase AffiniPure Goat Anti-Mouse IgG (H+L), Cat# 115-035-003; Peroxidase AffiniPure Goat Anti-Rabbit IgG (H+L), Cat# 111-035-003, both from Jackson ImmunoResearch; bovine anti-chicken IgY-HRP, Cat# sc-2917 from Santa Cruz Biotechnology.

### Secondary antibodies for IF

Chicken anti-Rabbit IgG (H+L) Cross-Adsorbed Secondary Antibody, Alexa Fluor 488, Cat# A21441; Goat anti-Mouse IgG (H+L) CrossAdsorbed Secondary Antibody, Alexa Fluor 546, Cat# A11003; Alexa Fluor™ Plus 647 Phalloidin, Cat#A30107; all from ThermoFisher Scientific; Abberior star 635P conjugated to phalloidin for STED imaging, Cat# ST635P-0100-20UG from Abberior.

### Cell culture and cell treatment

Patient-derived wild type and mutant fibroblast cells were isolated as described in [5] and cultured in MCDB medium supplemented with 5% fetal bovine serum, 1% Penicillin/Streptomycin, 1% Chemically Defined Lipid Concentrate, 1% HEPES, 1% GlutaMAX Supplement, 0.3% Insulin-Transferrin-Selenium, 1 ng/mL basic Fibroblast Growth Factor, 2 ng/mL Epidermal Growth Factor, 5 µg/mL Vitamin C, 250 nM hydrocortisone and 7.5 U/mL heparin. HEK293T cells used for lentiviral production were cultured and transfected with the appropriate plasmids as described in [34]. All cell culture was performed at 37°C in a humidified atmosphere containing 5% CO_2_. Wild type cells were treated with different concentrations of colchicine (Cat #C9754, Sigma) for 1 hour at 37°C as indicated in Figure 4A. Colchicine was dissolved in ethanol and ethanol was used to treat the “No colchicine” sample.

### Cell growth and cell migration assay

For cell growth assay, 20 000 cells were seeded in 24-well plates. Cell numbers were determined each day up to 96 hours by cell counting with trypan blue. For cell migration assays, 75 000 cells were plated in 24-well plates. 48 hours later the cells were pre-treated with 10 μg/ml mitomycin C (Cat# M5353, Sigma Aldrich) for 2 h to block proliferation and a cell-free area was created by scratching the monolayer with a 10-μl pipette tip. Subsequently, the cells were washed once with PBS. After that, MCDB medium, supplemented with 5% FBS and all the medium components listed above was placed on the top of the cells. Following scratching, images were recorded at several locations along the scratch line by using a ZOE Fluorescence Imager (BioRad) with a 20x objective. After 22 hours, images were acquired at the same locations. The area devoid of cells was determined by using the Fiji plugin MRI Wound Healing tool (Montpellier Resources Imagerie, CNRS). The area in percentage was calculated where the cells migrated in related to the original scratch area.

### Determination of F-actin and G-actin content

Samples of wild type and G302A mutant fibroblasts were lysed according to the protocol described in the G-actin / F-actin In Vivo Assay Kit (Cat# BK-037, Cytoskeleton Inc.). After a preclearing centrifugation step, G-actin and F-actin were separated by ultracentrifugation. The supernatant containing G-actin, and the pellet (resuspended in milliQ water supplemented with cytochalasin D) containing F-actin was loaded on an SDS-PAGE gel and the amount of actin was visualized by using a pan-actin antibody. After HRP-conjugated secondary antibody incubation, the membranes were incubated with chemiluminescence substrate and developed on Hyperfilms. Bands were quantified using ImageJ 1.53c.

### Plasmid

The active form (aa. 578-1198, i.e. the C-terminal part of the protein) of human mDia (DIAPH1 (NM_005219) Human Tagged ORF Clone, Origene, cat # RC216318) was subcloned into a lentiviral destination vector (Addgene, Cat# 17454), already containing an EGFP coding sequence, using gateway cloning. Lentiviral supernatants were generated by co-transfection of HEK293T cells with a three-vector lentiviral system: using the EGFP-mDia expression vector combined with the lentiviral packaging and envelope plasmids pRSV-Rev, pMDLg/pRRE and pCMV-VSV-G (kind gift of Prof. Guillaume Charras).

### Lenti-viral Packaging

Lenti-viral packaging of EGFP-mDia was performed using HEK293T cells. Transfection was carried out using polyethylenimine, branched (PEI, Sigma, Cat# 408727) as a transfection reagent, which was prepared according to [43]. 8.22·10^6^ HEK cells were seeded in T75 flask the day before transfection. The plasmid mixture contained 10 µg construct plasmid, 3.75 µg VSV-G, 1.875 µg Rev and 1.875 µg RRE virus plasmids. PEI was used in 1:2 ratio. HEK transfection medium was exchanged to cultured DMEM medium 3-4 hours after transfection. Viral supernatant was harvested after a total of 48 hours and filtered through 0.45 µm filter. The virus was concentrated with the Lenti-X Concentrator of TakaraBio (Cat# 631232) according to the manufacturer’s instruction. For transduction fibroblasts were trypsinized and cultured in 50% MCDB and 50% viral supernatant for 24 hours, supplemented with polybrene (Santa Cruz Biotechnology, Cat# sc-134220, 4 µg/ml final concentration). 1·10^5^ cells were seeded on a gelatine (0.5% in PBS)-coated 3.5 cm TPP dish.

### Monolayer stretching

Fibroblast monolayers were cultured on special chambers (CuriBio Inc., Cat# CS-2x25-UPF for 5 mm x 5 mm chambers, and Cat# CS-0144-UPF) used in the Cytostretcher LV instrument of CuriBio. The bottom of the chamber was treated with 0.2 mg/ml polydopamine solution (dissolved in 10 mM TRIS buffer, pH 8.5) for 2.5 hours, washed 3 times with sterile water to create a hydrophilic surface for gelatin coating. Stretching was carried out with 0.5%/sec velocity and the monolayer was kept stretched for 15 minutes before fixation. 2·10^4^ cells and 1.25·10^5^ were seeded in 5 mm x 5 mm and 12 mm x 12 mm chambers, respectively and cells were grown for 36 hours before stretch. Cells from 12 mm x 12 mm chambers were harvested at the indicated time points after stretch in 25 mM HEPES, pH 7.4, 150 mM NaCl, 1 mM EGTA, 1% NP-40, 10% glycerol, supplemented with the following protease and phosphatase inhibitors: 10 mM sodium pyrophosphate, 10 mM sodium fluoride, 5 mM sodium vanadate, 1 mM PMSF and cOmplete, EDTA-free protease inhibitor cocktail (Sigma, Tokyo, Japan). Lysates were centrifuged with 5000× *g* for 5 min at 4 °C and the supernatant was snap frozen for further immunoblotting.

### Immunofluorescence staining of fixed monolayers

Cells were fixed with Image-iT™ Fixative Solution (ThermoFisher Scientific, Cat# R37814) for 15 minutes. After that, cells were washed with HBSS, permeabilized (0.25% Triton X-100 in TBS-T, 10 min RT), blocked (1% BSA in TBS-T, 1 hour RT), and incubated with the primary antibody (dilutions prepared in 1% BSA-TBS-T for cofilin – 1:200, pFAK – 1:150, tubulin, 1:100, ZO-1 – 1:100; incubation was done overnight at 4 °C). After thorough washing in TBST, cells were stained simultaneously with the appropriate secondary antibodies (dilutions were prepared as 1:1000) and phalloidin (1:1000 in 1% BSA-TBST) for 1 hour at RT, washed in TBS-T and PBS. The cell nuclei were labeled with Hoechst for 10 minutes (Cat# 62249, ThermoFisher Scientific) and finally washed in PBS prior to imaging. The imaging was carried out in an antifade solution, 1% DABCO 33-LV (Sigma, Cat# 290734), containing 50% glycerol in PBS.

### Cell lysis and immunoblotting

Wild type and mutant fibroblasts were harvested in 25 mM HEPES, pH 7.4, 150 mM NaCl, 1 mM EGTA, 1% NP-40, 10% glycerol, supplemented with the following protease and phosphatase inhibitors: 10 mM sodium pyrophosphate, 10 mM sodium fluoride, 5 mM sodium vanadate, 1 mM PMSF and cOmplete, EDTA-free protease inhibitor cocktail (Sigma, Cat# 4693132001). Lysates were centrifuged with 5000 g for 5 minutes at 4°C and the supernatant was snap frozen for further immunoblotting. Proteins were separated using standard SDS-PAGE gel electrophoresis with 12% SDS-PAGE gels, transferred to PVDF membranes for immunoblot analysis using a wet blot transfer system. After blocking the membranes were stained with specific primary antibodies (overnight at 4 °C) as indicated in each figure. Next morning, the membranes were incubated with the appropriate HRP-conjugated secondary antibodies (1.5 hours at room temperature). Bands were visualized by using a chemiluminescence substrate and developed on Hyperfilms. Bands were quantified using ImageJ.

### Confocal and STED microscopy

Non-stretched and stretched samples were analyzed on a Nikon Ti2 confocal microscope. For confocal images of samples used in monolayer stretching pictures were taken by using a 20× lens (numerical aperture: 0.75, field of view 140 µm × 140 µm, resolution: 1000 × 1000 pixels). All other confocal images were taken by using a 60x lens (numerical aperture: 1.40, oil, field of view 120 µm × 120 µm, resolution: 1000 × 1000 pixels). STED images of fixed cells were taken by using a 100x lens (numerical aperture: 1.45, oil). Depletion was carried out with a 775 nm laser, with 20% laser power. Images were taken from randomly selected areas of the cell monolayer.

### Quantification of actin filament bundle width from the STED images

STED images were deconvoluted by using the Huygens Professional software 23.04, Scientific Volume Imaging BV. Quantification of actin width was carried out after the deconvolution by using a macro plugin of ImageJ, which is using a Gaussian fit of the line profile of phalloidin fluorescence. The width is given as the fitted value corresponding to the 2x standard deviation (2xSD) of the Gaussian distribution. All analyses were performed blinded, such as data analyzers were unaware of the genotypes.

### Quantification of the number and size of focal adhesions from the confocal images

The number of focal adhesions was determined in 140 × 140 μm area. The threshold was set between 0.9 and 1.2 % and the image was converted to a binary one. As a result, the focal adhesions appeared as white spots and the particles between sizes of 0.1 μm^2^ and infinity were counted using the Particle Analysis plugin in Fiji 1.53c software. Images were taken from randomly selected areas of the cell monolayer. All analyses were performed blinded, such as data analyzers were unaware of the genotypes.

### Quantification of cofilin relocalization upon stretch

First, confocal images measured in the z direction were summed by using the ImageJ “Maximal Intensity Projection” function. After that, stacked images of both phalloidin- and cofilin-stained samples were converted to a binary image by using a threshold between 80-90% to identify the areas having a fluorescence signal. Then, the Particle analysis tool of ImageJ was applied on the masked images (to identify the areas having “no” fluorescence signal and a size limitation of 20-100 μm^2^ was used to exclude tiny and very large holes) to determine the size of each “empty” area. After that, the corresponding “empty” areas of phalloidin- and cofilin-stained images were used for further calculations. We subtracted the area of the phalloidin-stained “holes” form the area of the cofilin-stained “holes” and divided by the area of the phalloidin-stained “holes” to get the ratio of the area where cofilin staining cannot be detected with the used threshold. Then, the numbers were converted to percentage for each condition (wild type or mutant, stretched or non-stretched) were plotted and compared. All analyses were performed blinded, such as data analyzers were unaware of the genotypes and treatments.

### AFM force spectroscopy of patient-derived fibroblasts

AFM imaging (Igor Pro 6.37 software) was performed in contact mode with an MFP3D AFM [44] using Bruker, MSCT-A probes (nominal typical spring constant = 70 pN/nm). Cantilevers were calibrated by the thermal method [45]. Cell monolayer samples were grown on circular microscope slides, which were mounted in the Bio-Heater module of the AFM. Imaging and force spectroscopy were carried out in a temperature-controlled liquid environment at 37°C. First an AFM image was taken, then *in situ* force spectroscopy was carried out collecting 100 force curves on selected 3x3 µm regions of cell surfaces (force mapping). The individual force curves were recorded with a vertical Z-piezo movement speed of 1 μm/s, until the force set point (0.5 nN) was reached, then the tip was retracted. Elastic moduli were obtained by fitting the indentation curves with the blunted pyramidal model as described in Rico *et al*. [46]. For the calculations [47] the irregular pyramid shape with a semi-included angle of 20°, and with a spherical cap radius of 10 nm was used as described for the Bruker, MSCT-A probe. Poisson ratios of the tip and the sample were set to 0.2 and 0.5, respectively.

### Statistical Analyses

Statistical analyses were carried out in GraphPad Prism 4.01. Significance was determined by one-way or two-way ANOVA followed by a Bonferroni post-test. Differences between groups were considered statistically significant if p < 0.05.

## AUTHOR CONTRIBUTIONS

É.G. and A.V. designed the experiments. K.D., É.G., T.B., K.P. and A.V. performed experiments. K.D., É.G., K.P. and A.V. analyzed data. K.D., É.G., K.P., N DD., M.K. and A.V. wrote the manuscript. All authors contributed feedback for the manuscript. All authors have read and agreed to the published version of the manuscript.

## FUNDING

This action has received funding from the ERA-NET COFUND /EJP COFUND Programme with co-funding from the European Union Horizon 2020 research and innovation programme, with the project No. 2019-2.1.7-ERA-NET-2020-0001 coordinated by the National Research, Development and Innovation Office (M.K. and A.V.). This research was also funded by the Thematic Excellence Programme (TKP2021-EGA-23) of the Ministry for Innovation and Technology in Hungary, within the framework of the Therapeutic Development and Bioimaging thematic programs of the Semmelweis University.

## ACKNOWLEDGEMENT

The authors thank the members of the PredACTINg consortium for helpful discussions. The technical contribution of Béláné Szénási from the Department of Molecular Biology of Semmelweis University with ultracentrifugation and the contribution of Ágnes Kovács with the western blot workflow are greatly acknowledged.

## DATA AVAILABILITY STATEMENT

The original contributions presented in the study are included in the article. Further inquiries can be directed to the corresponding author.

## CONFLICT OF INTEREST

The authors declare no conflicts of interest.

## LEGENDS TO THE FIGURES

**Figure S1.**
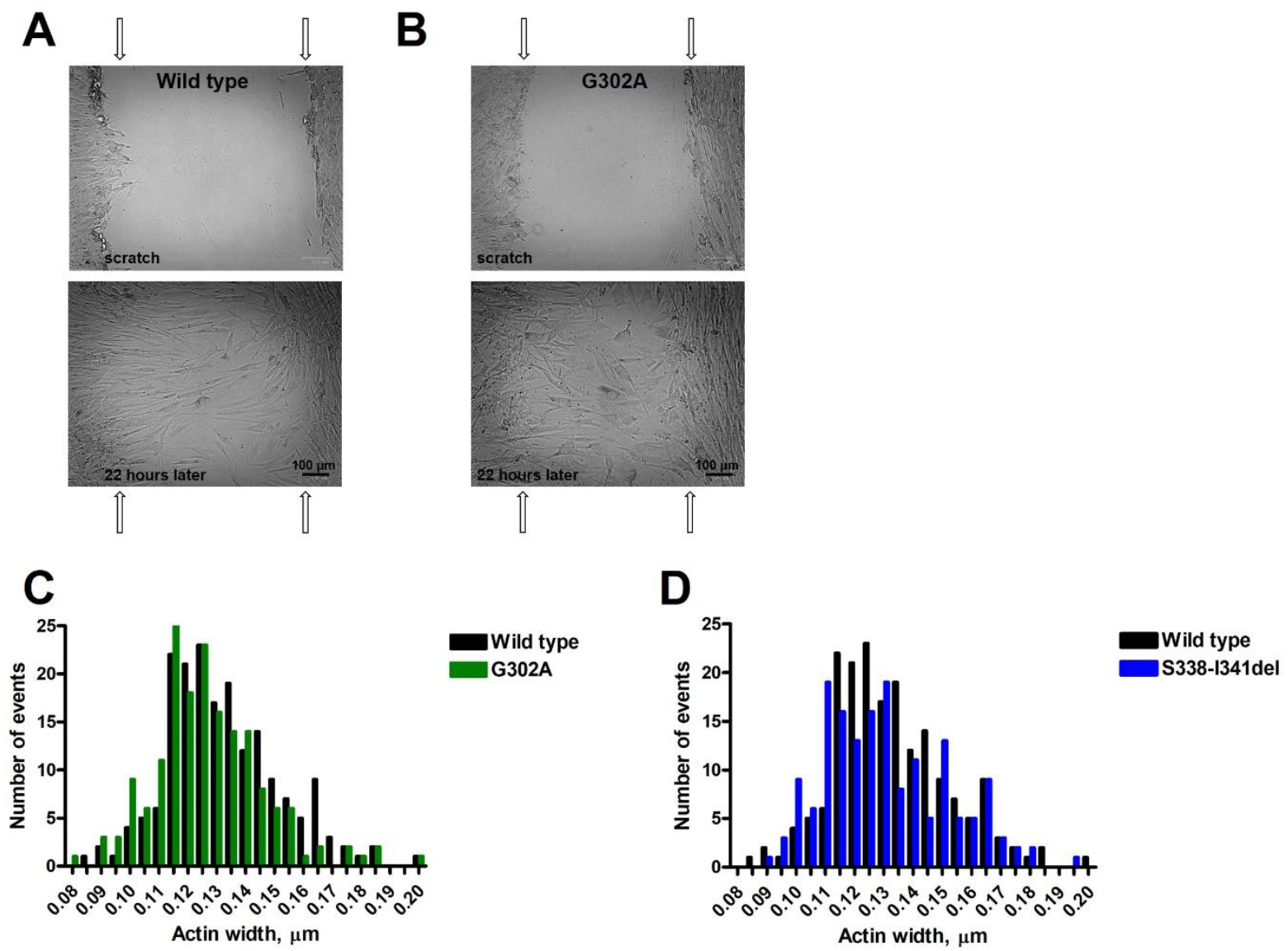
related to Figure 2. Fibroblast motility assayed by *in vitro* wound healing for wild type **(A)** and G302A mutant **(B)** fibroblast cells. Wound closure was photographed after 22 h. Arrows show the wound margins at time 0. Pairwise comparison of the histograms of actin width determined for wild type and G302A fibroblasts **(C)**, or wild type and S338-I341del fibroblasts **(D)**.

**Figure S2.**
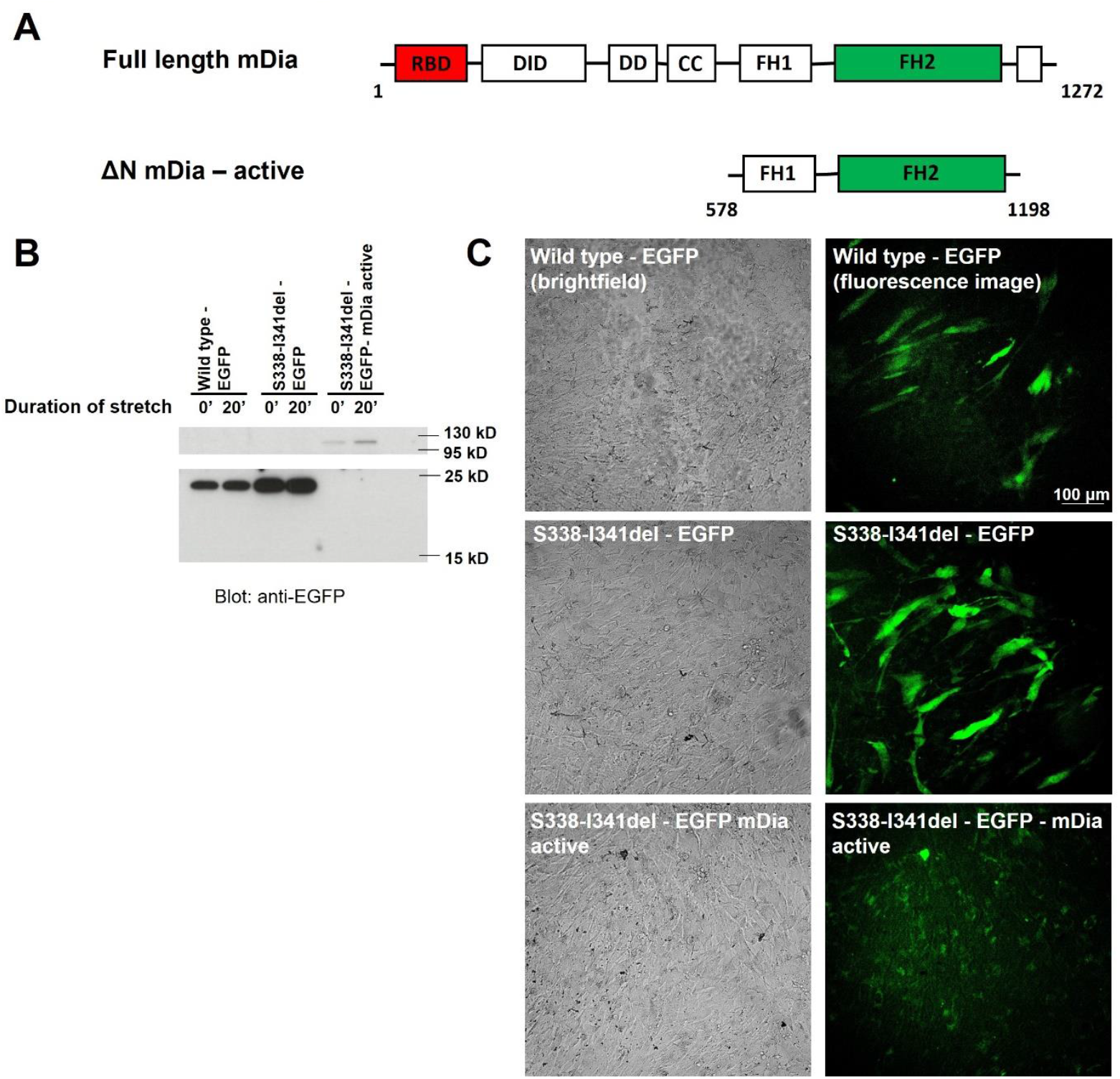
related to Figure 7. **(A)** Domain structure of full length and active mDia. Active mDia is a truncated form, where the N-terminal part of the full-length protein (containing the RBD, DID, DAD and CC domains) is missing (called also ΔN). **(B)** Western blot of the samples of the same experiment shown in Figure 7A, blotted for anti-EGFP. Molecular weight markers are shown to indicate the appropriate size of the molecules expressed in wild type and S338-I341del cells. **(C)** Examples of brightfield and the corresponding fluorescence image of each monolayer used in the experiment shown in Figure 7A are represented. EGFP expression was visualized with a 20x objective on the ZOE fluorescence cell imager (Bio-Rad).

**Figure S3.**
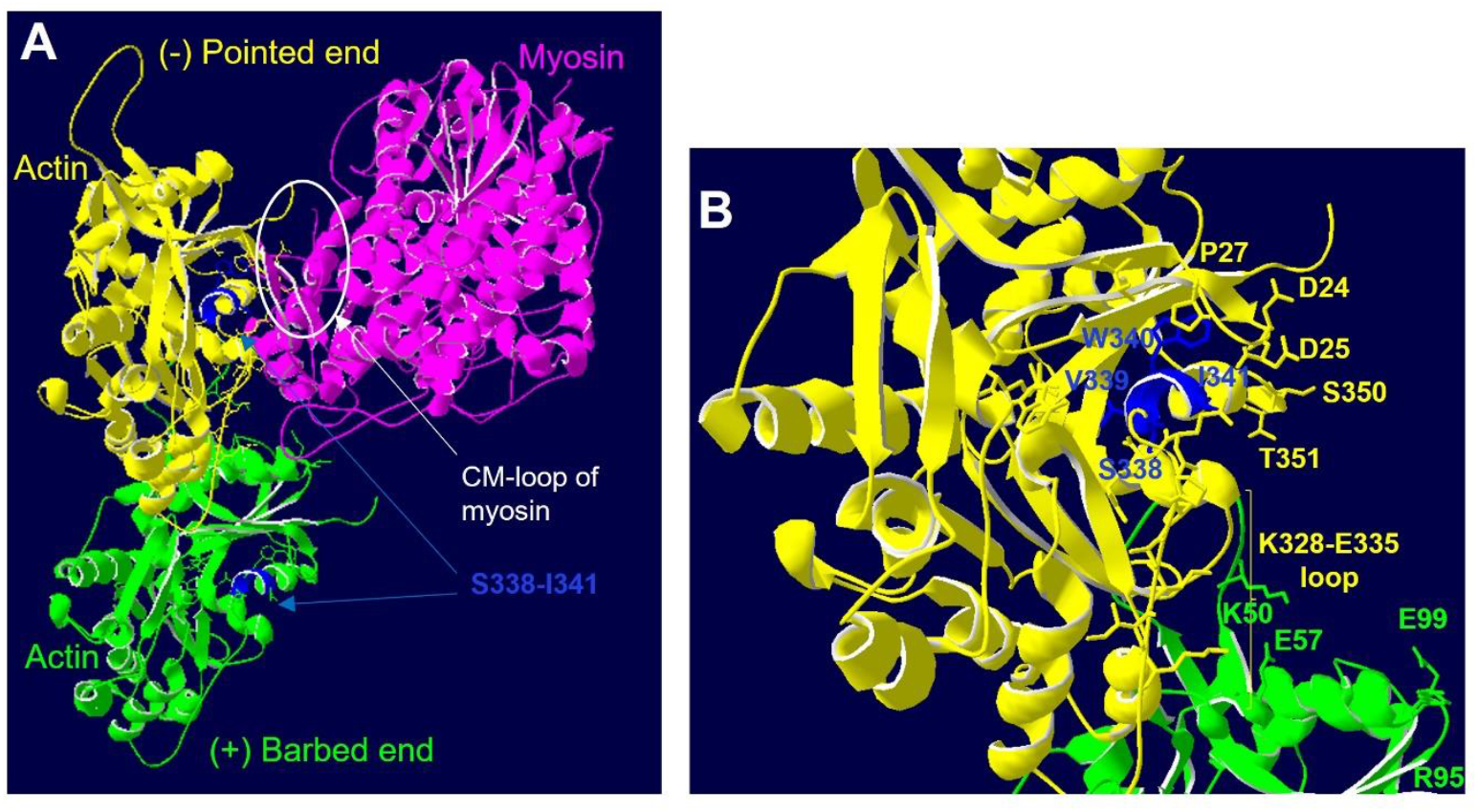
related to Figure 6. **(A)** Myosin (purple) binding to the actin filament as represented by the published cryo-EM structure (PDB code: 5JLH, [37]). Two actin protomers are shown within the filament, colored yellow and green, creating binding sites for myosin. The S338-I341 segment, which is the beginning of a helix located in SD1, is colored dark blue. Residues interacting with myosin are labeled as stick models. **(B)** Zoomed-in view of the surrounding of the helix having the segment of S338-I341, showing the stacking interaction between W340 (blue) and P27 (yellow) from the loop interacting with the cardiomyopathy (CM) loop of myosin.

## Notes

### Competing Interest Statement

The authors have declared no competing interest.

